# Coalescence of rhizobial communities in soil interacts with fertilization and determines the assembly of rhizobia in root nodules

**DOI:** 10.1101/2020.01.16.908863

**Authors:** Josep Ramoneda, Johannes Le Roux, Stefanie Stadelmann, Emmanuel Frossard, Beat Frey, Hannes Andres Gamper

**Author notes:** **Corresponding author:** Josep Ramoneda. **Competing interests** The authors declare no competing financial interests.

## Abstract

Soil microbial community coalescence, whereby entire microbial communities mix and compete in a new environmental setting, is a widespread phenomenon whose applicability for targeted root microbiome assembly has not been studied. Using a legume shrub adapted to nutrient poor soil, we tested for the first time how the assembly of communities of rhizobial root nodule symbionts is affected by the interaction of coalescence and fertilization. Seedlings of the rooibos [*Aspalathus linearis* (Burm.f.) Dahlg.], were raised in pairwise mixtures of soil from cultivated and uncultivated land of five farms, as well as the individual mixture components. A fragment of the symbiosis maker gene, *nodA*, was sequenced to characterize the taxonomic turnover of the rhizobia associated with all root nodules at the age of eight month. Soil mixing promoted taxonomic turnover in the rhizobial communities, while fertilization amplified such turnover by increasing the number of rhizobia that became more abundant after soil mixing. Soil mixing and fertilization had a synergistic effect on the abundance of a particular taxon, which was rare in the component soils but became highly abundant in fertilized plants raised in soil mixtures. These findings provide the first evidence that fertilizer addition can interact with soil microbial community coalescence, probably through increasing the chances for rare strains to prioritize root nodule colonization. The combination of soil mixing and fertilizer addition may be a still unexplored measure to (re)introduce root microbial mutualists in arable land.

## Introduction

The merging of entire microbial communities is considered an important driver of microbial diversity and community function, but has been largely overlooked in studies on microbial community assembly (Rillig et al., 2015). During these co-called community coalescence events, distinct microbial assemblages often interact under novel environmental settings (Rillig and Mansour, 2017). When this happens, the degree of co-evolution among the members of each community, i.e. the level of community cohesion, is expected to determine which community dominates in the novel assemblage (Tikhonov, 2016; Sierocinski et al., 2017; Lu et al., 2018). For example, this is the case of methanogenic archaeal communities, in which metabolic functions among the taxa from mixed communities have been coupled to maximize the efficiency by which available resources are taken up by the community as a whole (Sierocinski et al., 2017). Another scenario appears when coalescing microbial communities have low levels of cohesiveness: in this case, the individual fitness of the component taxa under novel environmental conditions will define the structure of the new mixed microbial assemblage (Rillig and Mansour, 2017).

Soil microbial communities are expected to undergo frequent mixing events (Rillig et al., 2016a). Such events, for example soil tillage in agricultural systems, can involve communities with high cohesiveness (e.g. microbial communities isolated in soil aggregates, Rillig et al., 2017), and low cohesiveness (e.g. arbuscular mycorrhizal fungal propagules within the soil matrix, Vályi et al., 2016). When it comes to plants that depend on mutualistic symbioses with microbes, soil manipulation through mixing can provide host plants with new and diverse soil microbial assemblages they can select from (Mueller and Sachs, 2015). In agricultural contexts, this may occur under dramatically altered abiotic conditions, e.g. elevated soil nutrient content (Hartman et al., 2018). While the mixing of soil microbial communities is thought of as a promising intervention to support plant production (Rillig et al., 2016b), we still lack basic knowledge on how plant-associated microbial communities respond to mixing and changes in soil abiotic conditions.

Nitrogen-fixing rhizobia (α- and β-Proteobacteria) represent a particularly interesting system where community composition in response to coalescence might be important, but has never been studied before. This is maybe surprising given the high relevance of rhizobium-legume interactions to plant production. Endosymbiotic rhizobia provide their legume hosts with fixed atmospheric nitrogen in exchange for photosynthetates. Rhizobia are unique in that they can only multiply endophytically, i.e. inside root nodules, where they depend on the transfer of carbohydrates synthesized by the host plant. Upon nodule decay, rhizobia are released into the soil, where they can survive saprotrophically (Denison and Kiers, 2011). This means the rhizobial assemblages found in the soil do not represent cohesive assemblages, but mixes of distinct populations derived from the surrounding legume species, which have evolved different degrees of specificity with them (Sprent et al., 2017). Thus, legumes are presented with new and more diverse rhizobia when soil communities are mixed (Rillig et al., 2017). This, in turn, increases the probability of legumes interacting with rhizobia that are more efficient at fixing or transferring nitrogen (i.e. a sampling effect of rhizobial diversity *sensu* Loreau and Hector, 2001).

Rhizobial community composition and diversity is most dependant on the availability of suitable plant hosts (Sprent et al., 2017), but also importantly on soil nitrogen and phosphorus contents and water availability (Vuong et al., 2017). Thus, novel rhizobial assemblages created by coalescence may respond to soil fertilization *a priori* in unpredictable ways. On the one hand, fertilization can decrease the selectivity of legumes when recruiting rhizobial partners (Kiers et al., 2003), thus allowing for any potential rhizobial partner to colonize root nodules. On the other hand, some rhizobia may be better adapted than others to elevated soil nutrient and water availability, and therefore have increased the chances to prioritize colonization of root nodules. Stochastic microbial community assembly processes, dominated by priority effects, are known to be stronger when nutrient availability high, leading to unpredictable community assemblages (Chase, 2010).

Here we addressed whether fertilization of coalesced rhizobial communities can favour the overdominance of particular rhizobial taxa in the root nodules of an agriculturally-important and symbiotically promiscuous legume, rooibos (*Aspalathus linearis*). We hypothesized that the mixing of different soils would allow for rhizobium strains uncommon in either of the soil components to colonize rooibos root nodules, thus creating a distinct symbiotic rhizobial community. Furthermore, we tested whether fertilization increased the likelihood for such over-dominant strains to emerge following coalescence. This would point at the potential of combining soil microbial community coalescence and fertilization as interventions to enhance sampling effects of microbial diversity in arable land (Bender et al., 2016).

We conducted a cross-factorial soil rhizobial community mixing experiment in which rhizosphere soils from cultivated (descendant) and wild (ancestral) rooibos populations were used separately and in 1:1 v/v mixes to grow rooibos seedlings for eight months. Half of these soils were amended with a constant amount of sheep dung (fertilizer) of known mineral nutrient composition. Rooibos represents an ideal system to test the effects of mineral nutrition and mixing of soil microbial communities because it grows on extremely nitrogen and phosphorus-poor soils (Hawkins et al., 2011), can associate with a wide range of rhizobial symbionts (Hassen et al., 2012; Le Roux et al., 2017), is initially grown in nurseries where soils can be manipulated, and because cultivated rooibos populations lie adjacent to its wild populations in the landscape. Thus, soil community coalescence events are also likely to occur under field conditions and could be of agronomic relevance.

## Materials and Methods

### Experimental design and setup

To examine the interactive effects of fertilization and soil mixing on the assembly of root-nodulating rhizobia in rooibos nodules, we set up a cross-factorial pot experiment using rooibos genotypes from the same commercial cultivar. Our experiment was designed with the factors i) ‘location’ with five levels [Blomfontein (Blo), Dobbelarskop (Dob), Landsklof (Lan), Matarakoppies (Mat) and Melkkraal (Mel)], each representing different geographical soil origins and landscape-scale replicates, ii) ‘soil origin’ with three levels, composed of soil from cultivated and adjacent uncultivated land from each geographical area and a 1:1 (v/v) volumetric mixture of those two soils, and iii) ‘fertilization’ with two levels, treatments with either the addition of sheep dung, or not. The experimental units were pots each containing one rooibos individual that were randomly arranged with respect to all of the 30 resulting experimental treatments (5 locations × 3 soil origins × 2 fertilization treatments) in ten spatial blocks. Each treatment was replicated ten times, leading to a total of 300 experimental units.

Soils used for the experiment were collected down to a depth of 30 cm from beneath cultivated and wild rooibos plants that were not more than 100 m apart, and locations that corresponded with different farming areas in the Suid Bokkeveld (Western Cape, South Africa). The locations were located between 5.5 and 33 km apart. Three sulfuric acid-scarified and surface-sterilized seeds of rooibos were placed approximately 1 cm below the soil surface in a line perpendicular to that formed by the two columns of sheep manure, as a way to prevent any direct contact between the roots and manure. Starting from one month post germination, only the first germinating and most vigorously growing seedling was kept by removing any of the two possible additional seedlings. Furthermore, other plants growing from seeds and root pieces of the field soils were removed periodically.

The experiment was initially setup in a polytunnel and, after five months of growth, plants were transferred to an outdoor shade house at the Agricultural Research Council Infruitec’s research station in Stellenbosch, South Africa.

The fertilization treatment consisted in the addition of 7.5 g air-dry dung from a nighttime enclosure (Afrikaans ‘kraal’) of sheep on one of the farms where soils were collected at the time of filling the pots. Sheep dung was found to increase the total N content in soils by up to 30% and the total P content by up to 50% (see Table S1 for the detailed mineral nutrient concentrations of the sheep dung and soils). Moreover, it was verified that no additional rhizobia were added by sequencing microbial communities in the fertilizer. The experiment ran from the beginning of August 2016 to the middle of April 2017.

### Root nodule harvest, DNA extraction and sequencing

At harvest, individual root systems were rinsed with pressurized water. Stems, leaves, and roots were separated and the total number of root nodules was counted and weighed after air-drying at room temperature for 1–3 hours before final storage on silica gel. All root nodules of each individual plant were pooled and milled using a 7 mm diameter glass bead in a 2 ml Eppendorf tube in a TissueLyser II (Qiagen, Hilden Germany). Total DNA was extracted from 25 mg of the resulting nodule powder using the NucleoSpin Plant II Kit (Macherey-Nagel, Düren, Germany) according to the manufacturer’s protocol with the sodium dodecyl sulfate-containing lysis buffer PL2 and elution in 100 μl 5mM Tris-EDTA (TE, pH 8.5) elution buffer (PE).

A symbiosis-specific rhizobial marker of ~455 bp-long (the nodulation gene *nodA*, encoding an N-acyltransferase of the rhizobial *nodA*BC operon), was amplified with the primers *NodA*univF145u (5’-TGGGCSGGNGCNAGRCCBGA-3’) and *NodA*Rbrad (5’-TCACARCTCKGGCCCGTTCCG-3’) (Moulin et al., 2001). PCR amplifications were performed in triplicates in final reaction volumes of 12.5 μl. Each reaction contained 3.75 μl PCR-grade water, 6.25 μl Multiplex PCR Master Mix (QIAGEN), 0.75 μl of each of the primers at a concentration of 10 μM and 1 μl of the template DNA. PCR reactions were run in DNA Engine Peltier Thermal Cyclers (Bio-Rad), using the following conditions: initial denaturation at 95°C for 5 min followed by 27 cycles of denaturation at 95°C for 30 s, primer annealing at 69°C for 90 s and primer extension at 72°C for 35 seconds, with a final extension at 68°C for 10 min. PCR products were kept at 10°C until storage at −20°C. The three amplification product replicates of each plant-root nodule extract were pooled and purified with home-made SPRI beads at a ratio of 0.8 times the reaction volume. The amplicon size and integrity was verified on the Agilent 2200 Tape Station (Agilent, USA) before dilution to 4 ng □1^−1^for equimolar pooling of all 266 samples for library preparation. The amplicons were 2×300 bp paired-end sequenced, using the Illumina MiSeq v2 chemistry and PhiX as internal standard at a concentration of 48.99% on an Illumina MiSeq sequencer at the Genetic Diversity Center of ETH Zürich, Switzerland.

### Bioinformatics of sequencing data

The 3’-ends of the *nodA* paired-end reads were trimmed with seqtk (https://github.com/lh3/seqtk) and ends were joined with FLASH (Magoč and Salzberg, 2011). The priming sites were removed with USEARCH (v10.0.240) as described in Edgar (2011). PRINSEQ-lite (Schmieder and Edwards, 2011) was used to check the quality of all sequences. The unique sequence reads were obtained with the dereplication command *fastx_uniques* in USEARCH and Illumina sequencing errors were corrected and chimeras removed in UNOISE3. This generated zero-radius operational taxonomic units (ZOTUs). These are expected to represent the true biological sequences and hence possibly different bacterial strains. To further exclude possible variants due to PCR and sequencing errors, we clustered the ZOTUs at 99% nucleotide sequence identity, using the USEARCH function *cluster_smallmem*. The count table containing the information about the read numbers and thus ZOTU abundance was extracted, using the USEARCH function *otutab*. Taxonomic assignment and a chimera check were done in SINTAX (Edgar, 2016), using all available *nodA* sequences from NCBI GenBank as the reference database (https://blast.ncbi.nlm.nih.gov/Blast.cgi). The count table was generated using the *otutab* command in USEARCH. The phylotaxonomic affiliations ofthe ZOTUs and a further chimera check were done in SINTAX (Edgar, 2016) in comparison to all *nodA* sequences on NCBI GenBank. Raw sequences have been deposited in Dryad (https://doi.org/10.5061/dryad.zgmsbcc6q).

### Statistical analysis

The ZOTU occurrence and prevalence per sample were corrected for unequal sequencing depth by intra-extrapolation to the median read number of 8720 per sample after exclusion of four samples whose read number was less than 200, using the *phyloseq_inext* function of the *iNEXT* package v.2.0.19 of R (Hsieh et al., 2016). The richness, diversity and compositional and structural divergence of the ZOTU communities were determined, based on these median-read adjusted ZOTU abundances in *phyloseq* package v1.16.2 and *vegan* package v2.5-5 of R (McMurdie et al., 2013; Oksanen et al., 2007). Square-root-transforming the ZOTU abundances prior to the α- and β-diversity analyses did not qualitatively change the results. Effects of the experimental treatments on alpha diversity measures were tested using a generalized mixed effects model: *Alpha diversity* ~ *Fertilization*Soil origin*, *random= ~1|Block/Location.* P-values for the results of the mixed effect models were obtained by running ANOVA on the model fits.

The effects of the experimental treatments on the composition and structure of the rhizobial communities in the root nodules were then visualized using non-metric multidimensional scaling (NMDS) of Bray-Curtis dissimilarities calculated with the R package *vegan* (v2.5-5, Oksanen et al., 2007). The effect size and significance of the study factors ‘location’, ‘soil origin’ and ‘fertilization’, and the ‘soil origin x fertilization’ interaction were determined by permutational multivariate analysis (PERMANOVA) of the variance of the Bray-Curtis dissimilarities with the *adonis* function of *vegan*, using 9999 permutations. The nested experimental design was controlled by including the argument *strata = Farm.* Pairwise PERMANOVA with the *pairwise.adonis* function of the *pairwiseAdonis* R package (Martinez-Arbizu, 2019), was used to compare the communities assembled from the pure soils (‘cultivated’ and ‘uncultivated’) with those assembled from the mixed soils. The percentage of variance explained by each experimental factor and levels of the factor ‘soil origin’, were determined based on the ratio of the factor- or factor level-specific and total sum of squares. Subsequently, to explore whether fertilization imposed a common abiotic filter to the rhizobial communities, we determined the mean pairwise distances between the centroids of communities from fertilized and unfertilized plants. This was done using the function *distance_between_centroids* of the package *usedist* (v0.1.0, https://cran.r-project.org/web/packages/usedist/usedist.pdf) on a Bray-Curtis dissimilarity matrix, following the formula of Apostol and Mnatsakanian, (2003).

To visualize and statistically support the interaction between the factors ‘soil origin’ and ‘fertilization’, and hence demonstrate the impact of fertilization on the coalescence of the communities of rhizobial symbionts, we ran a differential abundance analysis for the treatment pairs ‘mixed-uncultivated’ and ‘mixed-cultivated’, using the R package *DESeq2* (v1.24.0, Love et al., 2015). Briefly, the function *DESeq* computes a negative binomial generalized linear model (GLM) that compares the total number of reads belonging to each ZOTU in the treatments being compared (in this case the pairs Mixed-Uncultivated and Mixed-Cultivated). This was done after normalization of untransformed reads across samples with the mean read count over all samples and scaled with a size factor specific to each sample as described in Love et al., (2015). The output from this analysis is a list of all shared ZOTUs between treatments and the differential log2fold increase in abundance of each ZOTU in either treatment. Benjamini-Hochberg’s false discovery rate correction for multiple testing was used when determining the significance of the log2-fold changes in abundance of all the ZOTUs shared between two of the levels of the study factor ‘soil origin’ per fertilization level. Those ZOTUs that had statistically significant increases in abundance between soil treatments were counted, their log2fold increases averaged, and their means compared using a Welch 2-sample t-test. In this way, we could test whether fertilization affected the number of ZOTUs differentially abundant between soil treatment pairs (Mixed-Uncultivated and Mixed-Cultivated).

## Results

After quality filtering, a total of 931 254 *nodA* sequence reads were retained for further analyses. All 93 ZOTUs identified in the root nodules belonged to the genus *Mesorhizobium*. The average ZOTU richness per plant was high (34 ± 9 ZOTUs) and was affected by geographical location only (Table S2). The strongest explanatory factor for rhizobial community structure was the location from which the soils were initially collected (i.e. geography), followed by its interaction with soil origin (i.e. cultivated vs wild; Table 1; Fig. 1). From this, it was expected that any effects of soil mixing on rhizobial community structure would be found within soil location, and therefore locations were treated as independent rhizobial community types. Location alone explained almost six times more variation in community structure (24.6%) than soil origin alone (4.2%). Neither fertilization or its interaction with soil origin had any effects on rhizobial community structure (Table 1).

**Table 1.**
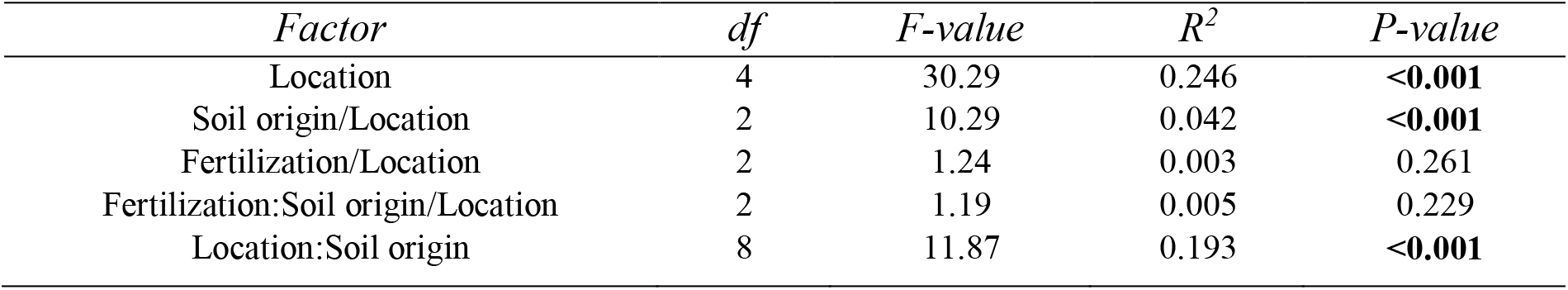
Permutational analysis of variance (PERMANOVA) on the effects of the experimental factors ‘location’, ‘soil origin’, ‘fertilization’ and their interactions on the structure of the symbiotic rhizobial communities of rooibos. [*Aspalathus linearis* (Burm.f.) Dahlg.]. The analysis was carried out on Bray-Curtis dissimilarities calculated on the relative abundance of 93 zero-radius operational taxonomic units (ZOTU) defined at 99% nucleotide sequence identity of a 455 bp-long fragment of the nodulation gene *nodA*.

**Figure 1.**
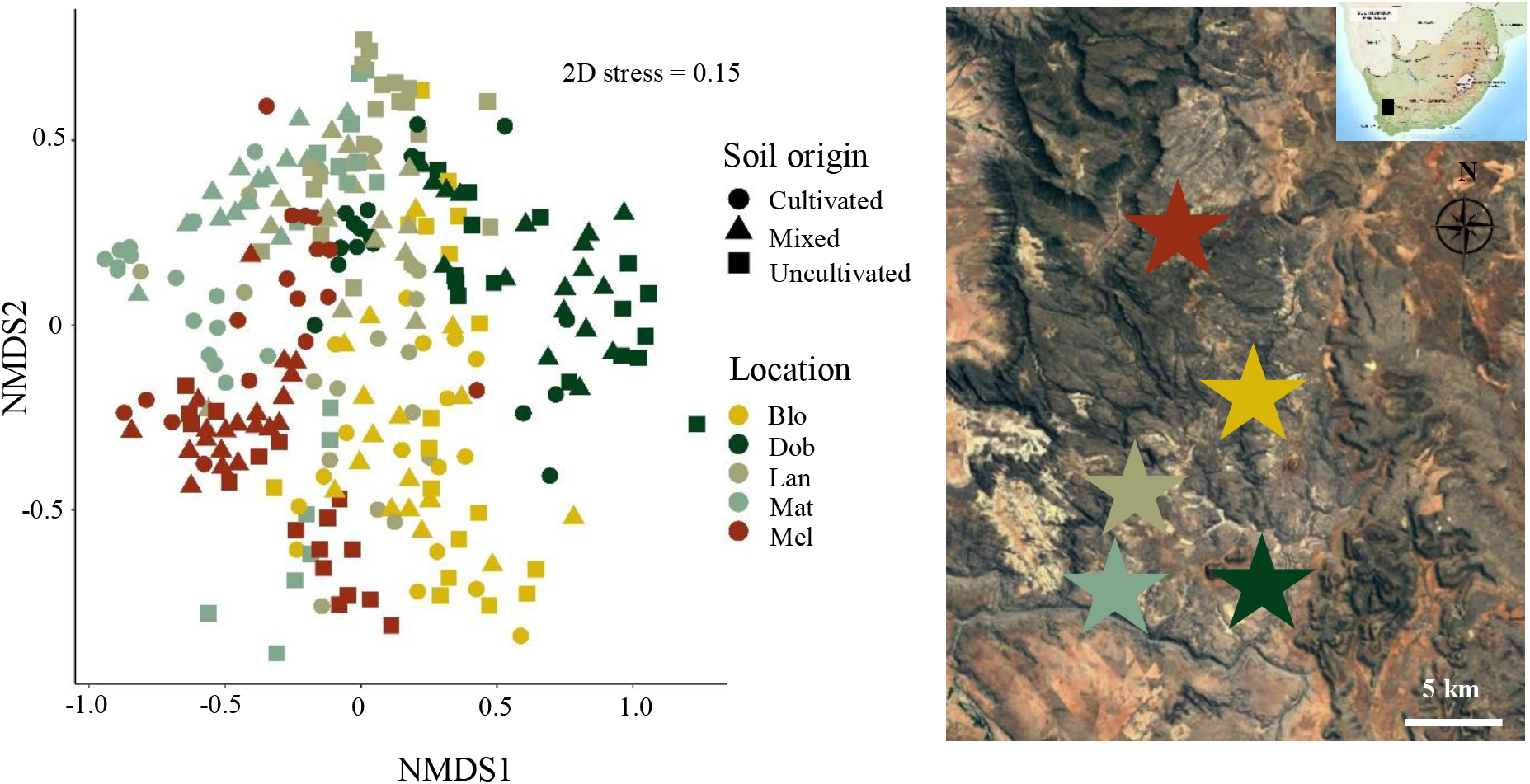
Non-metric multidimensional scaling (NMDS) plot of the symbiotic rhizobial communities of eight month-old rooibos [*Aspalathus linearis* (Burm.f.) Dahlg.] seedlings raised in soils from cultivated and uncultivated land or the volumetric 1:1 mixture of those from five locations. The analysis is based on Bray-Curtis dissimilarities calculated of the relative abundances of 93 zero-radius operational taxonomic units (ZOTU) defined at 99% nucleotide sequence identity of a 455 bp-long fragment of the nodulation gene nodA.

Within locations from where soils were collected, plants grown in cultivated and uncultivated soils always contained distinct rhizobial communities (Table 2). A large majority of plants grown in mixed soils harbored distinct rhizobial communities from those grown on non-mixed soils (i.e. cultivated or uncultivated), regardless of fertilization (Table 2). However, when comparing the magnitude of the differences between unfertilized and fertilized plants using distances between group centroids, three patterns emerged. Firstly, in communities from locations Blomfontein (Blo), Dobbelarskop (Dob), and Matarakoppies (Mat) fertilization led to both increases and decreases in rhizobial community similarity, depending on the soils being compared (Fig. 2A, B). Secondly, in the rhizobial communities from Landsklof (Lan), fertilization led to an increase in community similarity in all three soil type comparisons (i.e. fertilization filtered the rhizobial communities in a directional, homogenizing manner) (Fig. 2C). Thirdly, in communities from Melkkraal (Mel), fertilization promoted a decrease in the structural similarity between communities from different soil origins (Fig. 2D, E).

**Table 2.** Pairwise permutational analysis of variance of the symbiotic rhizobial communities of rooibos [*Aspalathus linearis* (Burm.f.) Dahlg.] fertilized or not with sheep manure. Rhizobial communities assembled from soils collected at five locations from adjacent ‘cultivated’ and ‘uncultivated’ land, and the volumetric 1:1 ‘mixture’ of those. The analysis was carried out on Bray-Curtis dissimilarities calculated of the relative abundance of 93 zero-radius operational taxonomic units (ZOTU) defined at 99% nucleotide sequence identity of a 455 bp-long fragment of the nodulation gene *nodA*. For each pair of soils the R^2^- values and significance levels are shown. Locations: Blomfontein (Blo), Dobbelarskop (Dob), Landsklof (Lan), Matarakoppies (Mat) and Melkkraal (Mel). P-values were Bonferroni-corrected for 5-times testing, corresponding to the five different locations.

| Fertilization | Soil | Locations |  |  |  |  |
| --- | --- | --- | --- | --- | --- | --- |
|  |  | Blo | Dob | Lan | Mat | Mel |
| Unfertilized | uncultivated-cultivated | 0.290*** | 0.390*** | 0.606*** | 0.416*** | 0.166<sup>ns</sup> |
|  | uncultivated-mixture | 0.177** | 0.331*** | 0.140<sup>ns</sup> | 0.448** | 0.189<sup>ns</sup> |
|  | cultivated-mixture | 0.121<sup>ns</sup> | 0.336*** | 0.476*** | 0.182<sup>ns</sup> | 0.129<sup>ns</sup> |
| Fertilized | uncultivated-cultivated | 0.227** | 0.456*** | 0.237*** | 0.548** | 0.327** |
|  | uncultivated-mixture | 0.218** | 0.234** | 0.029<sup>ns</sup> | 0.588*** | 0.305** |
|  | cultivated-mixture | 0.130<sup>ns</sup> | 0.403** | 0.205** | 0.459** | 0.221*** |

P > 0.05, ns; p < 0.05, \*; p < 0.01, \*\*; p < 0.001, \*\*\*

**Figure 2.**
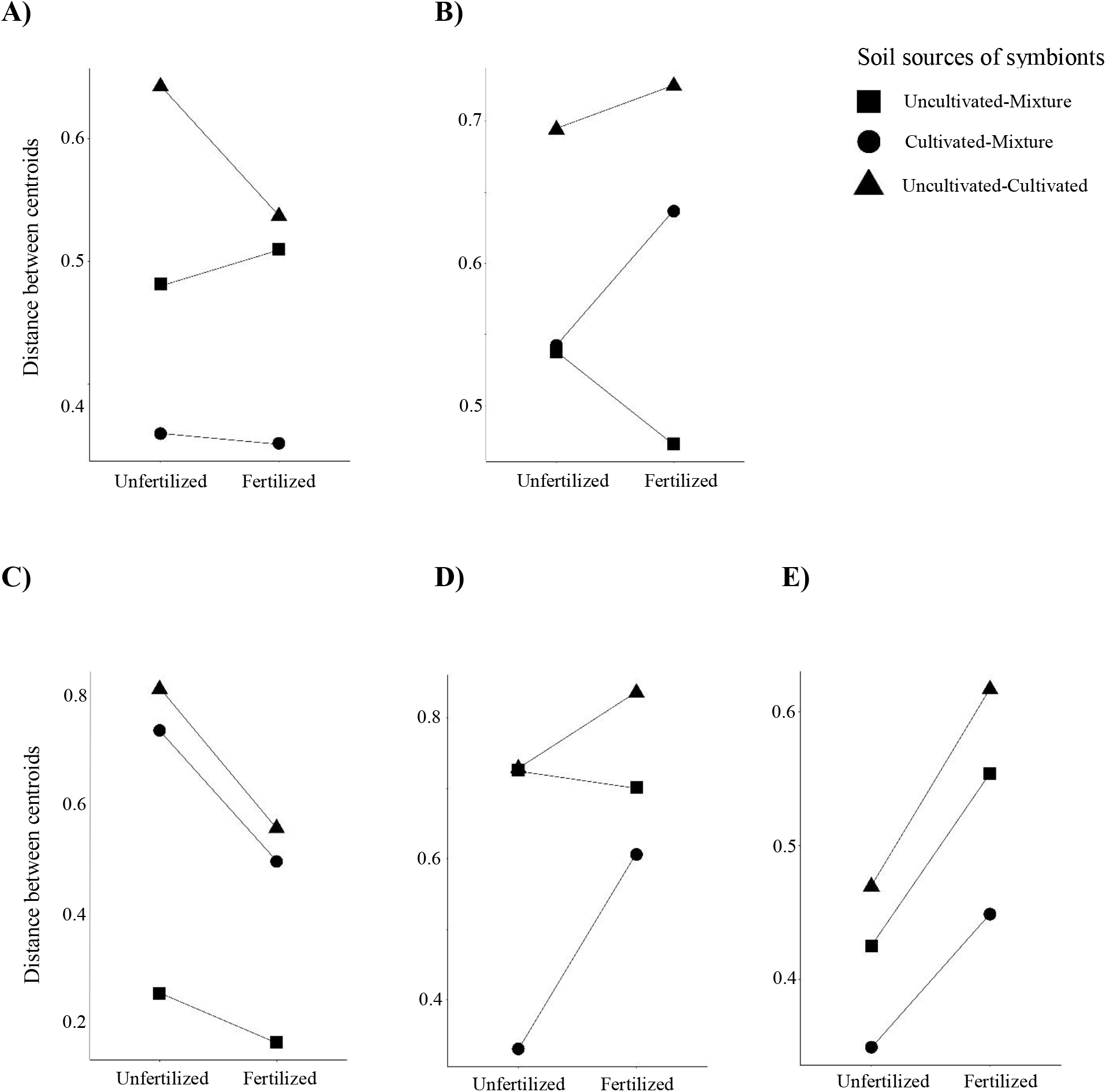
Differences between the symbiotic rhizobial communities of eight month-old rooibos [*Aspalathus linearis* (Burm.f.) Dahlg.] seedlings, assembled from soils from eight locations (A-E) and adjacent cultivated and uncultivated land, and the volumetric 1:1 mixture of those soils. These were either unfertilized or fertilized with sheep manure. The differences were calculated as the distances between the centroids of the communities of the non-metric multidimensional scaling plot of Figure 1. Changes in response to fertilisation are indicated as reaction norms: Lines pointing upwards point at divergence, lines point downwards indicate convergence and horizontal lines no change in community structuring. A: Blomfontein, B: Dobbelarskop, C: Landsklof, D: Matarakoppies, E: Melkkraal.

In order to explore the interaction between fertilization and soil mixing in more detail, we compared the dissimilarity between unfertilized and fertilized communities only from plants grown in soil mixes. Importantly, only the two communities increasing community dissimilarity after fertilization had a significant change in the structure of mixed soil communities after fertilization (Matarakoppies: R^2^ = 0.384, P = 0.003; Melkkraal: R^2^ = 0.133, P = 0.035). When looking at the effects of fertilization on communities from either cultivated or uncultivated soils, we found disparate results. Rhizobia from plants grown in cultivated soils were affected by fertilization only in Landsklof (R^2^ = 0.196; P = 0.024), whereas communities from plants grown in uncultivated soils were affected in Matarakoppies and Melkkraal (Community C: R^2^ = 0.210; P = 0.002; Community D: R^2^ = 0.204; P = 0.044).

Fertilization led to an increase of both positive and negative responders to soil mixing (Fig. 3A). Particularly, the number of ZOTUs responding positively to soil mixing increased in response to concurrent fertilization of the seedlings. At the same time, the average increase in the abundance of ZOTUs responding positively to soil mixing did not change significantly by fertilization (t = 1.41, P = 0.217; Fig. 3B), pointing at mostly a compositional rather than structural taxon-turnover in the symbiotic rhizobial communities. Fertilized plants had on average 16.5 differentially abundant ZOTUs across all soil sources (min.: 15 ZOTUs, max.: 20 ZOTUs), while unfertilized plants had on average only 6.75 differentially abundant ZOTUs (min.: 3, max.: 11; Fig. 3A, Table S3).

**Figure 3.**
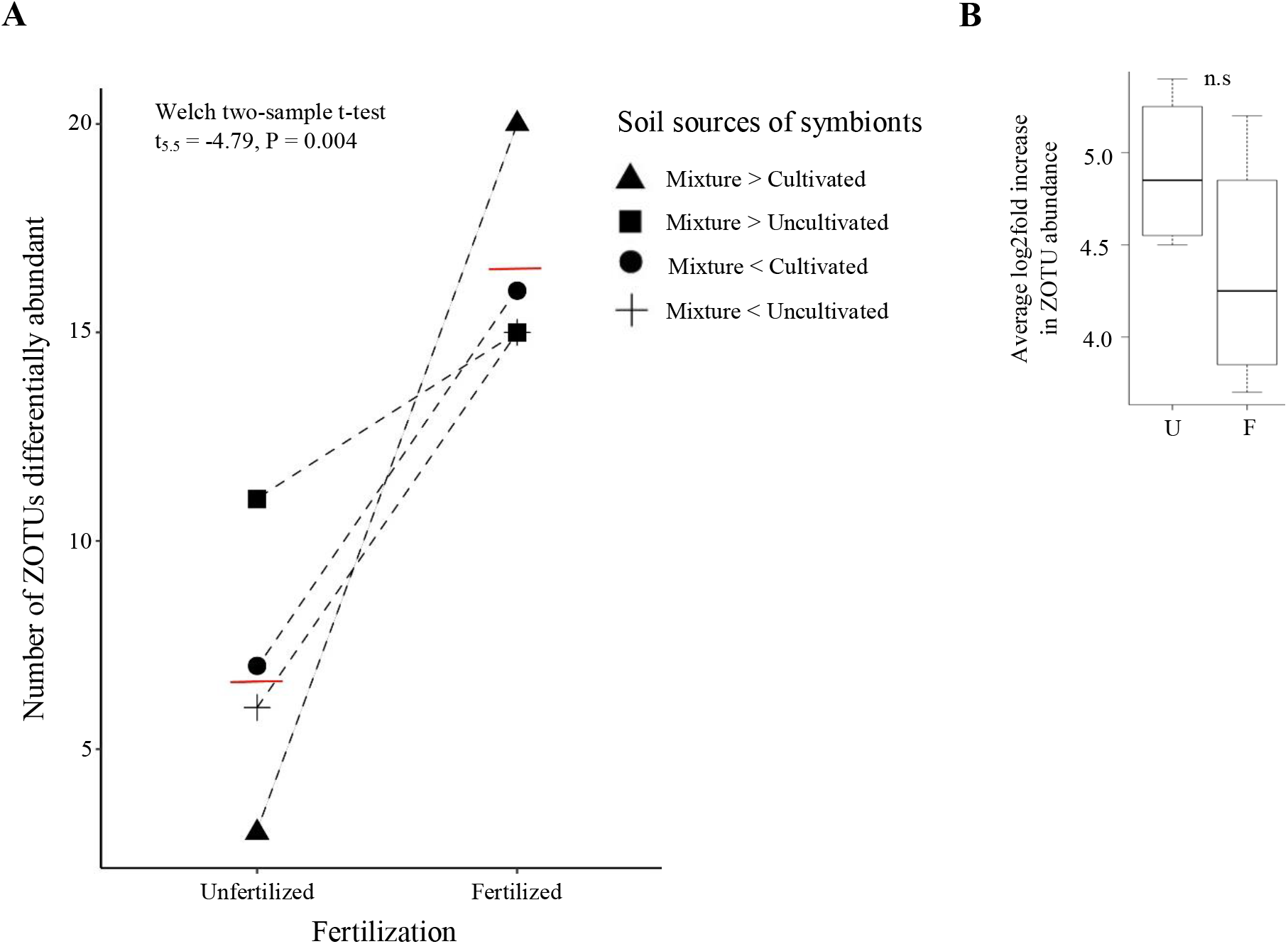
Effects of soil fertilization on the differential abundance of rhizobial ZOTUs in root nodules of rooibos plants grown on cultivated, uncultivated and 1:1 v/v mixes of these soils. A) Total number of rhizobial ZOTUs differentially abundant in root nodules of plants grown on cultivated and uncultivated soils compared to their 1:1 v/v mixes before and after fertilization. B) Average increase in rhizobial ZOTU read-normalized abundance in plants grown on unfertilized (U) and fertilized (F) soils. Statistically significant differences in ZOTU abundance were calculated with a generalized linear model (GLM) and differences between means were calculated using a Welch two-sample t-test with p ≤ 0.05 as a significance threshold. The differential fold-increase in rhizobial abundance is represented in a log2 scale.

Finally, we found ZOTU12 (*Mesorhizobium*) to have a large and significant increase in abundance for both the soil mixing and fertilization treatments (Fig. 4). Fertilization had a particularly remarkable effect; with this ZOTU having a 5.75-fold increase in read numbers, increasing from 1838 to 10570 sum-normalized reads, when unfertilized and fertilized mixed soils were compared (Fig. 4).

**Figure 4.**
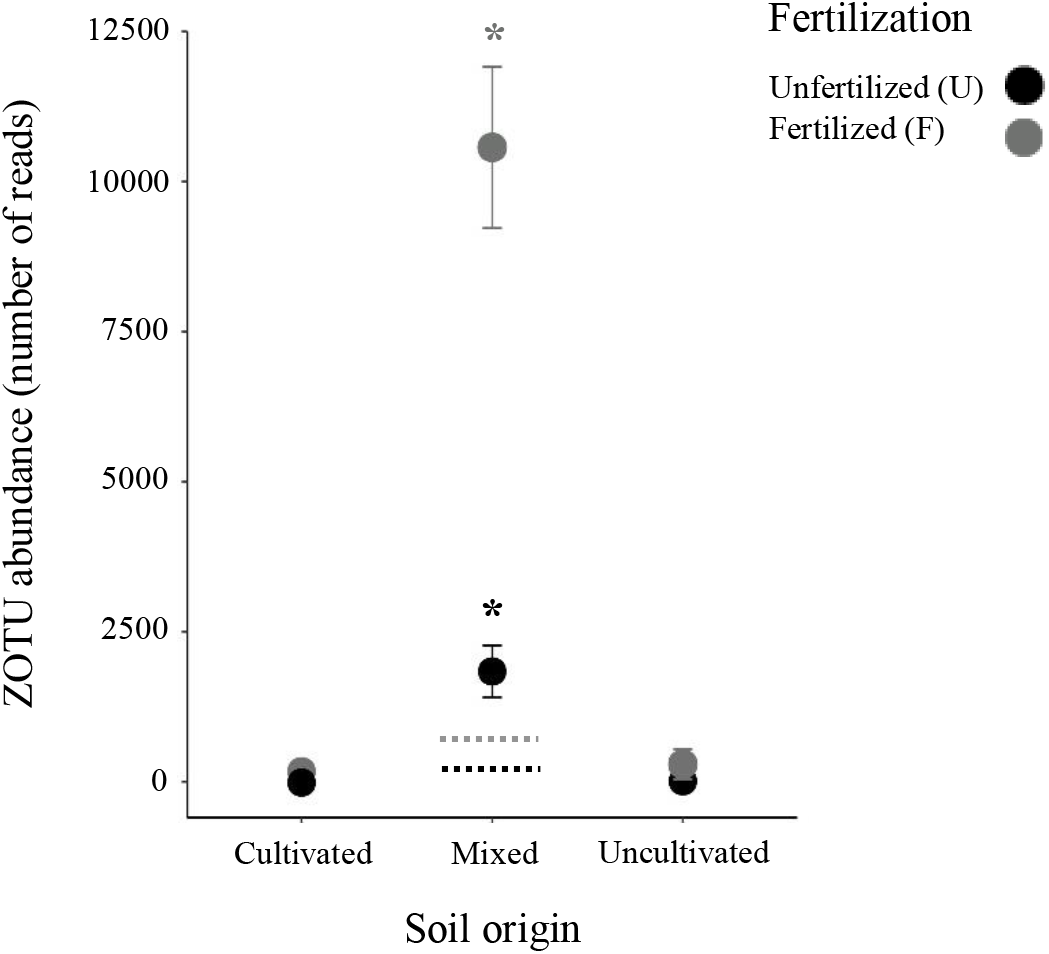
Abundance of one of 93 symbiotic zero-radius operational taxonomic units (ZOTU12) of the genus *Mesorhizobium* in eight month-old seedlings of rooibos [*Aspalathus linearis* (Burm.f.) Dahlg.]. Seedlings were raised in soils from cultivated and uncultivated land and 1:1 v/v mixes of those, from eight locations fertilized or not with sheep manure. The broken horizontal lines indicate the predicted abundance assuming additivity of the infection pressure in soil. The asterisks indicate that the abundances were significantly higher than predicted after mixing and fertilizing the seedlings, according to a statistical analysis with a negative binomial generalized linear model at the significance level of p ≤ 0.05.

## Discussion

Our findings indicate that soil nutrient content interacts with the structure of coalesced soil rhizobial communities, and can promote the dominance of otherwise rare rhizobial taxa. The effect of fertilization on the assembly of the communities of rhizobial symbionts was strongly location specific, both qualitatively as well as quantitatively. Depending on the geographical and ecological origin of the rhizobial source communities, the communities of rhizobial symbionts of rooibos diverged or converged (i.e. became more dissimilar or more similar to each other after fertilization). Consistent across all five locations, fertilization gave a greater number of ZOTUs a chance to become more abundant than when the seedlings were not fertilized, even though the average abundance of all these responsive ZOTUs in the communities of symbionts remained unchanged, as remained the richness and diversity of all ZOTUs. This means that fertilization gave a greater share of symbionts already becoming more abundant after soil mixing a chance to thrive even better. All in all, our experiment showed that there is synergism between community coalescence and fertilization. According to our knowledge, this is a novel finding for the symbiosis of legumes with rhizobia in general, relevant to the discussion of the effects of agriculture on bacterial communities and bacterial-root symbioses in particular (Bender et al., 2016).

Overall, soil nutrient content did not change the structure or diversity of root nodule rhizobial communities, indicating that fertilization did not impose a common filter to rhizobial survival. However, fertilization did allow for rhizobial communities from different soils to either become structurally more similar or different depending on their geographical origin. Communities consistently increasing in community similarity may have responded to a common environmental pressure imposed by fertilization. Elevated soil nitrogen, phosphorus, and organic matter are the main candidates to represent such common environmental filter, favouring only particular taxa (Vuong et al., 2017). Alterations to the availability of nutrients and water retention in soil must have acted mostly indirectly on the assembly of rhizobial root symbionts via changes to root growth and nutrient and carbohydrate allocation (Vuong et al., 2017). In particular, relief from phosphorus limitation after fertilization could have affected nutrient-efficient, slow-growing rhizobia and hence favored fast growing ones (Zahran, 1999). Although Zahran, (1999) discussed this at the level of rhizobial genera, it may also apply to the intrageneric or strain level, where there may exist more and less nutrient-conservative strains (Burghardt, 2019). Landsklof, the soil origin of the communities of rhizobial symbionts that all consistently became more similar in response to fertilization, was a location at which the physicochemical soil properties did not differ much between the cultivated and uncultivated land, except for soil pH, which was considerably higher under cultivation and also highest across all five locations (Table S1). It is thus well conceivable that adding manure to the soils had induced highly similar changes, and that the commonly observed acidification when ammonium is acquired by plants had reduced the difference in soil pH (Liu et al., 2017), explaining the consistency and convergence in the changes of the communities of rhizobial symbionts.

In contrast, we interpret that decreases in rhizobial community similarity may be the result of strong stochastic effects (e.g. inhibitory priority effects), whereby early colonization by one or a few rhizobium strains inhibit the establishment of others (Nemergut et al., 2013). Priority effects could have favored particular rhizobia that are not necessarily adapted to elevated soil nutrient contents, but that took advantage of the abundant soil and plant-derived resources available in order to prioritize root nodule colonization. In this way, pre-existing differences in rhizobial community composition between the different soils could have been amplified, making rhizobial community structure to shift in different directions depending on the soil origin. The effects of such stochasticity on community structure have been repeatedly observed for various taxonomic groups and communities. Previous research on freshwater mesocosms (Chase, 2007, 2010), microorganisms found in nectar and plant communities (Vannette and Fukami, 2014), and plant communities (Kardol et al., 2013), found that increasing nutrient supply enhances stochastic community assembly driven by priority effects, leading to increased community dissimilarity. For example, Vannette and Fukami (2014) showed that only when the sugar and aminoacid concentrations in nectar were high, did the first microbial strains to arrive and establish show strong dominance over later arriving strains.

Soil mixing promoted taxonomic turnover in the rhizobial communities, and fertilizer addition amplified this turnover by making a larger number of strains more abundant. Whether different strains had particular traits for fast colonization of root space, to obtain plant resources more efficiently, or to withstand nutrient-rich conditions in the soil, is still unclear. In rhizobia, the plant is normally the strongest filter to root nodule colonization, either by selecting which rhizobial taxa it associates with, or by differentially rewarding rhizobia already established in the nodules (Denison and Kiers, 2011). Plants, in the presence of abundant nutrients, rely less on effective symbiosis with rhizobia for nitrogen and this relax their pre-infection filtering of symbionts (Weese et al., 2015). This can allow for a wider range of strains, irrespective of their effectiveness, to prioritize nodule colonization, which would support the theory that bacterial traits would underlie the overdominance of a larger number of strains in the fertilized treatment (Barrett et al., 2015). At the same time, under fertilization a better nourished plant could provide more constant and abundant photosynthates, enhancing also the stochastic effects of root nodule community assembly (i.e. all strains would be equivalent in their ability to obtain plant photosynthetates, which would be non-limiting).

The fact that fertilization only increased the likelihood, but not the magnitude, of differential rhizobial abundance from mixed soils further supports the notion that higher soil nutrient content enhances the stochastic assembly of root nodule bacteria. This is supported by the fact that the relative abundance of all these responsive rhizobia to soil mixing and fertilization did not correlate with plant biomass production or foliar N contents (Table S4). Here, a tradeoff seems to exist as fertilization allowed for more different taxa to dominate in different plants, but with slightly lower average abundance across treatments. Under no fertilization, the few strains that differed between soil treatments became more abundant, were specifically rewarded by the host plant and/or survived under those soil conditions (Vuong et al., 2017). Importantly, one ZOTU was highly responsive to the combination of soil mixing and fertilization. This strain was rare in both cultivated and uncultivated soils and increased dramatically when soil mixtures were fertilized. Upon fertilization, this strain only increased its abundance in mixed soils, implying it may be not particularly well-adapted to high soil nutrient content, but rather to prioritize root nodule colonization. To our knowledge, there are no reports of rhizobia that consistently benefit more under coalescence, or of strains that are particularly good at recovering following soil disturbance.

Under agricultural conditions, soil microbial communities are usually biased towards few abundant taxa (Hartman et al., 2018). Coalescing communities from different locations could help to break over-dominances of taxa and allow subdominant taxa to become more abundant or to even reintroduce lost taxa. For example, in crops with extant populations of wild relatives, like rooibos (Hawkins et al., 2011), this could recover symbiotic interactions lost by crop domestication, breeding and intensive cultivation (Pérez-Jaramillo et al., 2017). Understanding aspects of stochastic root microbiome assembly is fundamental to our ability to manipulate microbial diversity and its possible function in plant production.

To our knowledge, this is the first study that assessed the combined effect of community coalescence and fertilization on the assembly of rhizobial symbionts of a crop plant. The finding that one symbiont particularly profited from combined fertilization and soil mixing is interesting, and possibly relevant for application. Inoculation with entire consortia increasingly becomes an agronomic measure in efforts to boost the growth and health of crop plants (Toju et al., 2018). Hence, measures to maintain and utilize rhizobial diversity under conditions of arable cropping are a subject that should receive more attention. This pot trial with actual field soil, suggests that it should be possible with simple measures to manipulate the communities of rhizobial symbionts of crop plants at the nursery stage, before transfer to plantations.

## Acknowledgements

This study was funded by the Mercator Research Program of the World Food Systems Center of ETH Zurich. Data produced and analyzed in this paper were generated in collaboration with the Genetic Diversity Center of ETH Zurich (GDC) and the Functional Genomics Center of the University of Zurich (FGCZ). JR acknowledges the assistance from Dr. Jean-Claude Walser, Dr. Andrea Patrignani and Dr. Weihong Qi. JLR acknowledges funding from Macquarie University’s Faculty of Science and Engineering and Department of Biological Sciences. Finally, the authors acknowledge the assistance of Noel Oettlé, Dr. Cecilia Bester, and the community of rooibos farmers in the Suid Bokkeveld for making the sampling in South Africa possible.

## Competing interests

The authors declare no competing financial interests.

## Notes

https://doi.org/10.5061/dryad.zgmsbcc6q

