## Supplementary Material for "Coalescence of rhizobial communities in soil interacts with fertilization and determines the assembly of rhizobia in root nodules"

**Table S1.** Physicochemical properties of soils from five plantations and adjacent wild populations of rooibos collected from cultivated and wild rooibos populations and sheep dung fertilizer. Locations/farms: Blo, Blomfontein; Dob, Dobbelarskop; Lan, Landsklof; Mat, Matarakoppies; Mel, Melkkraal.

| *Soil properties* | *Fertilizer (sheep dung)* | *Cultivated soil (plantations)*  *(Locations/farms)* | | | | | *Uncultivated soil (wild populations)*  *(Locations/farms)* | | | | |
| --- | --- | --- | --- | --- | --- | --- | --- | --- | --- | --- | --- |
|  |  | *Blo* | *Dob* | *Lan* | *Mat* | *Mel* | *Blo* | *Dob* | *Lan* | *Mat* | *Mel* |
| CEC (cmol kg^-1^) | - | 1.42 | 1.30 | 1.54 | 1.43 | 1.76 | 1.75 | 1.85 | 1.28 | 1.60 | 1.42 |
| WHC (% at 10 kPa) | - | 13.79 | 15.76 | 12.09 | 17.23 | 20.04 | 17.55 | 21.53 | 16.81 | 15.60 | 19.85 |
| Clay (%) | - | 7 | 5 | 5 | 5 | 9 | 7 | 7 | 5 | 7 | 7 |
| pH (CaCl_2_) | - | 5.78 | 5.76 | 6.15 | 5.53 | 5.55 | 5.89 | 5.48 | 5.51 | 5.60 | 5.29 |
| Organic C (%) | - | 0.54 | 0.65 | 0.54 | 0.75 | 1.11 | 1.21 | 1.79 | 0.87 | 3.93 | 1.05 |
| N (g kg^-1^) | 15.93 | 0.11 | 0.13 | 0.11 | 0.14 | 0.22 | 0.26 | 0.37 | 0.15 | 0.16 | 0.22 |
| P (g kg^-1^) | 1.86 | 0.020 | 0.001 | 0.003 | 0.010 | 0.007 | 0.005 | 0.030 | 0.007 | 0.010 | 0.020 |
| K (g kg^-1^) | 8.70 | 0.38 | 0.53 | 0.24 | 0.54 | 1.22 | 0.57 | 1.22 | 1.01 | 0.82 | 0.81 |
| δ^15^N (‰) | 16.36 | 20.40 | 19.75 | 23.68 | 22.29 | 14.52 | 10.87 | 10.31 | 20.26 | 19.31 | 13.91 |
| N:P ratio | 9.53 | 4.98 | 10.94 | 36.17 | 13.91 | 33.06 | 47.84 | 12.74 | 20.80 | 16.07 | 10.91 |
| Ca (g kg^-1^) | 12.30 | 0.06 | 0.11 | 0.06 | 0.05 | 0.12 | 0.14 | 0.16 | 0.06 | 0.06 | 0.08 |
| Mg (g kg^-1^) | 7.55 | 0.10 | 0.17 | 0.12 | 0.07 | 0.29 | 0.17 | 0.35 | 0.12 | 0.07 | 0.15 |

**Table S2**. Results of mixed model analyses on the effects of fertilization, soil origin and their interaction on the richness and diversity of rhizobia in root nodules of rooibos. The root nodules were analysed using gyrB and nodA DNA sequences, which was used to calculate zero-sums operational taxonomic units (ZOTU) based on 99% sequence identity. Richness represents the number of ZOTUs and diversity the Simpson’s diversity index. The experimental factor ‘fertilization’ involved sheep dung addition to soil, and ‘soil origin’ comprises the use of soils from five pairs of plantations and adjacent wild populations of rooibos and a 1:1 (v:v) mixture of those. The five locations (farms) were treated as a random and nested factor within fertilization and soil origin. There were 10 pot replicates per experimental treatment.

| *Factor* | *Richness* | | | |
| --- | --- | --- | --- | --- |
|  | *Richness* | | *Diversity* | |
|  | *F-value* | *P-value* | *F-value* | *P-value* |
| Location | 9.65 | **< 0.001** | 2.84 | **0.025** |
| Fertilization/Location | 0.75 | 0.387 | 3.10 | 0.080 |
| Soil origin/Location | 0.49 | 0.613 | 2.32 | 0.101 |
| Fertilization:Soil origin/Location | 0.45 | 0.638 | 1.74 | 0.178 |

**Table S3**. Summary of the differential abundance analysis depicting the fold-change in abundance of all ZOTUs shared between fertilized and unfertilized plants grown on mixed soils and plants grown on either cultivated or uncultivated soil. Log2-fold change in read numbers, standard error (SE) and Bonferroni-corrected P-values are provided. Statistically significant (P ≤ 0.05) fold-changes in ZOTU abundance are highlighted in bold. All ZOTUs belong to the genus *Mesorhizobium*.

**Unfertilized (Mixed vs Cultivated)**

| *Taxa* | *log2Fold Change* | *SE* | *P-value* |
| --- | --- | --- | --- |
| **ZOTU6** | **-7.805** | **1.073** | **0.000** |
| **ZOTU50** | **-7.223** | **1.333** | **0.000** |
| **ZOTU26** | **-6.191** | **1.601** | **0.001** |
| **ZOTU155** | **-4.710** | **0.858** | **0.000** |
| **ZOTU121** | **-4.565** | **0.904** | **0.000** |
| ZOTU66 | -4.175 | 1.991 | 0.162 |
| **ZOTU76** | **-4.100** | **1.271** | **0.010** |
| ZOTU106 | -3.749 | 1.447 | 0.064 |
| **ZOTU111** | **-2.899** | **0.785** | **0.002** |
| ZOTU18 | -2.335 | 0.984 | 0.099 |
| ZOTU28 | -2.179 | 0.896 | 0.092 |
| ZOTU139 | -2.159 | 1.242 | 0.256 |
| ZOTU49 | -2.155 | 1.037 | 0.162 |
| ZOTU102 | -2.069 | 1.030 | 0.172 |
| ZOTU91 | -1.914 | 1.913 | 0.598 |
| ZOTU132 | -1.899 | 1.952 | 0.603 |
| ZOTU70 | -1.870 | 1.992 | 0.619 |
| ZOTU159 | -1.746 | 2.063 | 0.659 |
| ZOTU99 | -1.737 | 0.961 | 0.234 |
| ZOTU79 | -1.490 | 1.498 | 0.598 |
| ZOTU114 | -1.479 | 1.412 | 0.598 |
| ZOTU37 | -1.445 | 0.718 | 0.172 |
| ZOTU109 | -1.402 | 0.820 | 0.256 |
| ZOTU48 | -1.296 | 1.201 | 0.598 |
| ZOTU173 | -1.273 | 0.875 | 0.395 |
| ZOTU36 | -1.258 | 1.236 | 0.598 |
| ZOTU172 | -1.189 | 1.568 | 0.700 |
| ZOTU21 | -1.140 | 1.632 | 0.722 |
| ZOTU14 | -1.067 | 1.180 | 0.637 |
| ZOTU148 | -0.923 | 1.427 | 0.756 |
| ZOTU123 | -0.862 | 1.498 | 0.778 |
| ZOTU112 | -0.761 | 1.818 | 0.880 |
| ZOTU97 | -0.648 | 0.760 | 0.659 |
| ZOTU174 | -0.632 | 0.838 | 0.700 |
| ZOTU120 | -0.629 | 1.047 | 0.769 |
| ZOTU162 | -0.594 | 1.139 | 0.799 |
| ZOTU40 | -0.511 | 1.828 | 0.936 |
| ZOTU130 | -0.400 | 1.447 | 0.936 |
| ZOTU145 | -0.369 | 1.517 | 0.951 |
| ZOTU136 | -0.361 | 1.740 | 0.963 |
| ZOTU62 | -0.134 | 1.138 | 0.987 |
| ZOTU82 | -0.127 | 2.698 | NA |
| ZOTU108 | -0.097 | 3.590 | 1.000 |
| ZOTU116 | -0.079 | 3.590 | NA |
| ZOTU150 | -0.075 | 2.052 | NA |
| ZOTU115 | -0.063 | 1.383 | 1.000 |
| ZOTU165 | -0.058 | 2.034 | NA |
| ZOTU100 | -0.043 | 3.225 | NA |
| ZOTU113 | -0.025 | 3.590 | NA |
| ZOTU154 | -0.005 | 2.324 | 1.000 |
| ZOTU45 | -0.004 | 2.668 | 1.000 |
| ZOTU110 | 0.000 | 0.000 | NA |
| ZOTU22 | 0.000 | 0.012 | 1.000 |
| ZOTU92 | 0.036 | 3.590 | NA |
| ZOTU101 | 0.090 | 3.590 | NA |
| ZOTU166 | 0.099 | 1.302 | 1.000 |
| ZOTU42 | 0.196 | 1.263 | 0.969 |
| ZOTU87 | 0.206 | 3.590 | NA |
| ZOTU68 | 0.215 | 3.590 | NA |
| ZOTU72 | 0.228 | 3.040 | NA |
| ZOTU86 | 0.341 | 1.087 | 0.933 |
| ZOTU90 | 0.345 | 2.029 | 0.969 |
| ZOTU71 | 0.453 | 2.983 | NA |
| ZOTU61 | 0.453 | 2.310 | 0.963 |
| ZOTU64 | 0.520 | 1.497 | 0.917 |
| ZOTU167 | 0.645 | 1.626 | 0.886 |
| ZOTU73 | 0.802 | 1.121 | 0.721 |
| ZOTU31 | 0.850 | 1.384 | 0.769 |
| ZOTU164 | 0.954 | 0.880 | 0.598 |
| ZOTU170 | 0.955 | 0.832 | 0.591 |
| ZOTU17 | 0.979 | 0.942 | 0.598 |
| ZOTU69 | 1.149 | 2.103 | 0.791 |
| ZOTU27 | 1.151 | 0.810 | 0.405 |
| ZOTU98 | 1.381 | 1.677 | 0.665 |
| ZOTU93 | 1.422 | 1.355 | 0.598 |
| ZOTU12 | 1.665 | 0.975 | 0.256 |
| ZOTU149 | 1.986 | 1.063 | 0.215 |
| ZOTU34 | 2.008 | 1.719 | 0.591 |
| ZOTU7 | 2.096 | 0.988 | 0.162 |
| ZOTU43 | 2.455 | 1.306 | 0.215 |
| ZOTU52 | 3.249 | 2.629 | 0.545 |
| **ZOTU19** | **3.660** | **1.181** | **0.014** |
| **ZOTU23** | **3.700** | **1.001** | **0.002** |
| ZOTU105 | 3.844 | 2.315 | 0.272 |
| **ZOTU33** | **5.989** | **1.462** | **0.001** |
| ZOTU63 | 6.417 | 2.815 | 0.118 |

**Unfertilized (Mixed vs Uncultivated)**

| *Taxa* | *log2Fold Change* | *SE* | *P-value* |
| --- | --- | --- | --- |
| **ZOTU21** | **-7.709** | **1.589** | **0.000** |
| **ZOTU76** | **-5.975** | **1.240** | **0.000** |
| **ZOTU43** | **-4.066** | **1.275** | **0.008** |
| **ZOTU14** | **-4.019** | **1.146** | **0.003** |
| ZOTU136 | -3.815 | 1.696 | 0.085 |
| ZOTU63 | -3.608 | 2.738 | 0.347 |
| **ZOTU62** | **-3.577** | **1.102** | **0.007** |
| ZOTU79 | -3.383 | 1.459 | 0.077 |
| **ZOTU99** | **-2.476** | **0.939** | **0.038** |
| ZOTU162 | -2.467 | 1.107 | 0.085 |
| ZOTU167 | -2.329 | 1.585 | 0.283 |
| ZOTU42 | -2.184 | 1.234 | 0.184 |
| ZOTU102 | -2.055 | 1.007 | 0.124 |
| ZOTU73 | -2.033 | 1.089 | 0.159 |
| ZOTU106 | -2.012 | 1.420 | 0.304 |
| ZOTU149 | -1.966 | 1.027 | 0.149 |
| ZOTU45 | -1.882 | 2.606 | 0.651 |
| ZOTU91 | -1.861 | 1.870 | 0.502 |
| ZOTU28 | -1.768 | 0.876 | 0.125 |
| ZOTU31 | -1.718 | 1.346 | 0.359 |
| ZOTU115 | -1.569 | 1.349 | NA |
| ZOTU173 | -1.552 | 0.855 | 0.172 |
| ZOTU139 | -1.545 | 1.217 | 0.359 |
| ZOTU165 | -1.403 | 1.986 | NA |
| ZOTU174 | -1.269 | 0.818 | 0.248 |
| ZOTU121 | -1.068 | 0.897 | 0.401 |
| ZOTU17 | -1.004 | 0.913 | 0.454 |
| ZOTU120 | -0.888 | 1.023 | 0.578 |
| ZOTU159 | -0.886 | 2.016 | 0.792 |
| ZOTU26 | -0.827 | 1.575 | 0.754 |
| ZOTU130 | -0.727 | 1.413 | 0.754 |
| ZOTU50 | -0.696 | 1.317 | 0.754 |
| ZOTU97 | -0.688 | 0.743 | 0.542 |
| ZOTU101 | -0.634 | 3.507 | NA |
| ZOTU92 | -0.634 | 3.507 | NA |
| ZOTU82 | -0.620 | 2.636 | NA |
| ZOTU90 | -0.581 | 1.981 | 0.879 |
| ZOTU113 | -0.490 | 3.507 | NA |
| ZOTU100 | -0.486 | 3.151 | NA |
| ZOTU108 | -0.333 | 3.507 | 0.982 |
| ZOTU69 | -0.292 | 2.054 | 0.968 |
| ZOTU132 | -0.289 | 1.909 | 0.968 |
| ZOTU87 | -0.267 | 3.507 | NA |
| ZOTU72 | -0.255 | 2.969 | NA |
| ZOTU68 | -0.191 | 3.507 | NA |
| ZOTU61 | -0.163 | 2.256 | 0.982 |
| ZOTU150 | -0.127 | 2.004 | NA |
| ZOTU93 | -0.086 | 1.321 | 0.982 |
| ZOTU66 | -0.082 | 1.951 | 0.982 |
| ZOTU49 | -0.040 | 1.020 | 0.982 |
| ZOTU22 | 0.000 | 0.011 | 1.000 |
| ZOTU110 | 0.000 | 0.000 | NA |
| ZOTU116 | 0.000 | 3.507 | NA |
| ZOTU71 | 0.052 | 2.914 | NA |
| ZOTU37 | 0.286 | 0.710 | 0.798 |
| ZOTU154 | 0.336 | 2.270 | 0.968 |
| ZOTU7 | 0.411 | 0.966 | 0.792 |
| ZOTU111 | 0.617 | 0.775 | 0.602 |
| ZOTU27 | 0.637 | 0.791 | 0.602 |
| ZOTU48 | 0.774 | 1.182 | 0.683 |
| ZOTU98 | 0.850 | 1.637 | 0.754 |
| ZOTU86 | 1.057 | 1.064 | 0.502 |
| ZOTU52 | 1.218 | 2.567 | 0.775 |
| ZOTU112 | 1.221 | 1.778 | 0.669 |
| ZOTU172 | 1.270 | 1.535 | 0.600 |
| ZOTU170 | 1.349 | 0.816 | 0.228 |
| ZOTU18 | 1.550 | 0.975 | 0.237 |
| ZOTU109 | 1.556 | 0.815 | 0.149 |
| ZOTU6 | 1.730 | 1.059 | 0.230 |
| ZOTU34 | 1.740 | 1.679 | 0.491 |
| ZOTU123 | 1.931 | 1.467 | 0.347 |
| ZOTU164 | 1.934 | 0.863 | 0.085 |
| ZOTU114 | 2.232 | 1.390 | 0.236 |
| **ZOTU155** | **3.188** | **0.844** | **0.001** |
| **ZOTU23** | **3.456** | **0.979** | **0.003** |
| ZOTU145 | 3.463 | 1.490 | 0.077 |
| **ZOTU12** | **3.698** | **0.961** | **0.001** |
| **ZOTU148** | **3.919** | **1.404** | **0.027** |
| **ZOTU64** | **4.073** | **1.468** | **0.027** |
| ZOTU70 | 4.200 | 1.956 | 0.099 |
| **ZOTU40** | **4.617** | **1.796** | **0.043** |
| **ZOTU36** | **5.366** | **1.221** | **0.000** |
| **ZOTU19** | **6.243** | **1.156** | **0.000** |
| **ZOTU166** | **6.323** | **1.286** | **0.000** |
| **ZOTU33** | **7.187** | **1.430** | **0.000** |
| **ZOTU105** | **7.559** | **2.264** | **0.006** |

**Fertilized (Mixed vs Cultivated)**

| *Taxa* | *log2Fold Change* | *SE* | *P-value* |
| --- | --- | --- | --- |
| **ZOTU26** | **-10.249** | **1.648** | **6E-09** |
| **ZOTU71** | **-5.456** | **2.283** | **0.037** |
| **ZOTU98** | **-4.986** | **0.951** | **0.000** |
| **ZOTU155** | **-4.189** | **0.719** | **0.000** |
| **ZOTU76** | **-4.135** | **0.975** | **0.000** |
| **ZOTU82** | **-3.968** | **1.633** | **0.034** |
| **ZOTU120** | **-3.907** | **0.722** | **0.000** |
| **ZOTU136** | **-3.644** | **1.564** | **0.042** |
| **ZOTU97** | **-3.513** | **0.558** | **0.000** |
| **ZOTU106** | **-3.486** | **1.008** | **0.002** |
| **ZOTU64** | **-3.451** | **1.145** | **0.007** |
| **ZOTU70** | **-3.327** | **1.314** | **0.027** |
| **ZOTU114** | **-2.885** | **0.979** | **0.009** |
| **ZOTU162** | **-2.703** | **0.810** | **0.003** |
| ZOTU115 | -2.698 | 1.396 | 0.097 |
| **ZOTU37** | **-2.395** | **0.645** | **0.001** |
| ZOTU123 | -2.321 | 1.432 | 0.175 |
| ZOTU112 | -2.281 | 1.480 | 0.197 |
| ZOTU86 | -2.187 | 0.993 | 0.055 |
| **ZOTU121** | **-1.974** | **0.680** | **0.010** |
| ZOTU154 | -1.789 | 1.661 | 0.377 |
| ZOTU92 | -1.672 | 3.525 | NA |
| ZOTU42 | -1.648 | 0.975 | 0.155 |
| ZOTU100 | -1.619 | 2.443 | NA |
| ZOTU132 | -1.550 | 1.493 | NA |
| ZOTU31 | -1.124 | 1.228 | 0.466 |
| ZOTU174 | -1.074 | 0.585 | 0.116 |
| ZOTU66 | -1.047 | 1.557 | 0.588 |
| ZOTU159 | -0.890 | 2.923 | NA |
| ZOTU48 | -0.858 | 0.998 | 0.496 |
| ZOTU167 | -0.728 | 1.189 | 0.614 |
| ZOTU165 | -0.622 | 1.590 | 0.746 |
| ZOTU108 | -0.527 | 3.017 | NA |
| ZOTU172 | -0.395 | 1.529 | 0.841 |
| ZOTU72 | -0.074 | 2.251 | NA |
| ZOTU107 | 0.000 | 3.528 | NA |
| ZOTU7 | 0.024 | 0.803 | 0.976 |
| ZOTU139 | 0.048 | 0.851 | 0.968 |
| ZOTU62 | 0.072 | 0.943 | 0.965 |
| ZOTU94 | 0.115 | 3.528 | NA |
| ZOTU113 | 0.121 | 3.528 | NA |
| ZOTU18 | 0.169 | 0.759 | 0.858 |
| ZOTU96 | 0.195 | 3.528 | NA |
| ZOTU111 | 0.309 | 0.597 | 0.672 |
| ZOTU163 | 0.316 | 3.528 | NA |
| ZOTU116 | 0.337 | 3.528 | NA |
| ZOTU101 | 0.395 | 3.527 | NA |
| ZOTU22 | 0.416 | 0.425 | 0.431 |
| ZOTU170 | 0.432 | 0.526 | 0.515 |
| ZOTU99 | 0.480 | 0.754 | 0.605 |
| ZOTU27 | 0.508 | 0.449 | 0.352 |
| ZOTU50 | 0.522 | 1.048 | 0.672 |
| ZOTU105 | 0.761 | 1.502 | 0.672 |
| ZOTU109 | 1.019 | 0.537 | 0.104 |
| ZOTU150 | 1.095 | 1.400 | 0.534 |
| ZOTU87 | 1.096 | 3.526 | NA |
| ZOTU73 | 1.196 | 0.597 | 0.084 |
| ZOTU90 | 1.227 | 1.789 | 0.587 |
| ZOTU79 | 1.290 | 1.017 | 0.295 |
| ZOTU102 | 1.298 | 0.902 | 0.230 |
| ZOTU130 | 1.323 | 0.939 | 0.238 |
| ZOTU149 | 1.423 | 0.701 | 0.081 |
| ZOTU91 | 1.533 | 1.355 | 0.352 |
| ZOTU145 | 1.898 | 1.236 | 0.197 |
| **ZOTU173** | **1.936** | **0.596** | **0.004** |
| ZOTU45 | 2.054 | 2.844 | 0.569 |
| ZOTU52 | 2.213 | 1.771 | 0.299 |
| **ZOTU23** | **2.248** | **0.843** | **0.019** |
| **ZOTU148** | **2.573** | **1.145** | **0.050** |
| **ZOTU6** | **2.632** | **1.079** | **0.034** |
| **ZOTU17** | **2.862** | **0.922** | **0.006** |
| **ZOTU164** | **2.954** | **0.722** | **0.000** |
| **ZOTU166** | **2.984** | **1.089** | **0.016** |
| ZOTU69 | 3.178 | 2.345 | 0.258 |
| **ZOTU36** | **3.200** | **1.046** | **0.007** |
| **ZOTU49** | **3.236** | **0.762** | **0.000** |
| **ZOTU14** | **3.334** | **0.920** | **0.001** |
| **ZOTU61** | **3.690** | **1.645** | **0.050** |
| **ZOTU34** | **3.904** | **1.230** | **0.005** |
| **ZOTU33** | **3.976** | **1.008** | **0.000** |
| ZOTU63 | 4.192 | 2.742 | 0.197 |
| **ZOTU93** | **4.239** | **1.126** | **0.001** |
| **ZOTU43** | **4.498** | **1.125** | **0.000** |
| **ZOTU12** | **5.393** | **0.844** | **4E-09** |
| **ZOTU68** | **6.223** | **1.920** | **4E-03** |
| **ZOTU40** | **6.740** | **1.622** | **2E-04** |
| **ZOTU28** | **6.849** | **0.828** | **5E-15** |
| **ZOTU21** | **6.949** | **1.101** | **4E-09** |
| **ZOTU19** | **11.534** | **1.003** | **1E-28** |

**Fertilized (Mixed vs Uncultivated)**

| *Taxa* | *log2Fold Change* | *SE* | *P-value* |
| --- | --- | --- | --- |
| **ZOTU62** | **-6.089** | **0.940** | **2E-09** |
| **ZOTU136** | **-5.140** | **1.573** | **0.005** |
| **ZOTU21** | **-5.119** | **1.091** | **0.000** |
| **ZOTU43** | **-3.959** | **1.129** | **0.002** |
| **ZOTU99** | **-3.898** | **0.759** | **0.000** |
| **ZOTU76** | **-3.691** | **0.983** | **0.001** |
| **ZOTU114** | **-3.436** | **0.989** | **0.002** |
| **ZOTU17** | **-3.422** | **0.911** | **0.001** |
| **ZOTU115** | **-3.375** | **1.407** | **0.046** |
| **ZOTU120** | **-3.295** | **0.734** | **0.000** |
| **ZOTU162** | **-3.028** | **0.817** | **0.001** |
| **ZOTU31** | **-3.020** | **1.235** | **0.044** |
| **ZOTU106** | **-2.977** | **1.020** | **0.014** |
| ZOTU90 | -2.766 | 1.790 | 0.247 |
| **ZOTU174** | **-2.728** | **0.588** | **0.000** |
| **ZOTU42** | **-2.686** | **0.981** | **0.021** |
| ZOTU26 | -2.618 | 1.668 | 0.241 |
| **ZOTU102** | **-2.366** | **0.905** | **0.028** |
| **ZOTU86** | **-2.352** | **0.999** | **0.050** |
| ZOTU154 | -1.945 | 1.673 | 0.427 |
| ZOTU69 | -1.871 | 2.349 | 0.632 |
| ZOTU132 | -1.753 | 1.504 | 0.427 |
| ZOTU48 | -1.684 | 1.007 | 0.210 |
| ZOTU164 | -1.566 | 0.722 | 0.074 |
| ZOTU93 | -1.518 | 1.114 | 0.342 |
| ZOTU108 | -1.360 | 3.035 | 0.826 |
| ZOTU109 | -1.235 | 0.536 | 0.055 |
| ZOTU92 | -1.146 | 3.549 | 0.841 |
| ZOTU73 | -1.127 | 0.586 | 0.131 |
| ZOTU101 | -1.120 | 3.547 | 0.841 |
| ZOTU159 | -1.113 | 2.942 | 0.826 |
| ZOTU163 | -1.070 | 3.548 | 0.841 |
| ZOTU14 | -1.070 | 0.915 | 0.427 |
| ZOTU22 | -1.038 | 0.428 | 0.045 |
| ZOTU107 | -1.031 | 3.549 | 0.841 |
| ZOTU96 | -1.019 | 3.549 | 0.841 |
| ZOTU94 | -1.014 | 3.549 | 0.841 |
| ZOTU111 | -0.998 | 0.602 | 0.210 |
| ZOTU100 | -0.993 | 2.461 | 0.826 |
| ZOTU165 | -0.972 | 1.601 | 0.737 |
| ZOTU113 | -0.917 | 3.549 | 0.854 |
| ZOTU72 | -0.911 | 2.265 | 0.826 |
| ZOTU149 | -0.869 | 0.697 | 0.402 |
| ZOTU37 | -0.836 | 0.651 | 0.385 |
| ZOTU123 | -0.817 | 1.450 | 0.750 |
| ZOTU91 | -0.816 | 1.361 | 0.737 |
| ZOTU116 | -0.781 | 3.548 | 0.865 |
| ZOTU49 | -0.676 | 0.748 | 0.562 |
| ZOTU139 | -0.671 | 0.860 | 0.635 |
| ZOTU82 | -0.641 | 1.658 | 0.826 |
| ZOTU98 | -0.575 | 0.974 | 0.737 |
| ZOTU173 | -0.351 | 0.590 | 0.737 |
| ZOTU18 | -0.338 | 0.767 | 0.826 |
| ZOTU79 | -0.240 | 1.019 | 0.862 |
| ZOTU63 | -0.168 | 2.751 | 0.962 |
| ZOTU87 | -0.140 | 3.547 | 0.968 |
| ZOTU150 | 0.088 | 1.408 | 0.962 |
| ZOTU172 | 0.213 | 1.543 | 0.921 |
| ZOTU167 | 0.470 | 1.206 | 0.826 |
| ZOTU130 | 0.505 | 0.944 | 0.765 |
| ZOTU28 | 0.506 | 0.832 | 0.737 |
| ZOTU121 | 0.644 | 0.709 | 0.562 |
| ZOTU97 | 0.700 | 0.568 | 0.404 |
| ZOTU27 | 0.794 | 0.452 | 0.185 |
| ZOTU170 | 0.893 | 0.542 | 0.210 |
| ZOTU7 | 0.912 | 0.808 | 0.444 |
| ZOTU34 | 0.928 | 1.228 | 0.646 |
| ZOTU64 | 0.962 | 1.174 | 0.622 |
| ZOTU145 | 1.297 | 1.244 | 0.490 |
| ZOTU66 | 1.431 | 1.578 | 0.562 |
| ZOTU23 | 1.947 | 0.848 | 0.055 |
| ZOTU71 | 2.584 | 2.314 | 0.444 |
| ZOTU112 | 2.607 | 1.505 | 0.190 |
| ZOTU45 | 2.836 | 2.864 | 0.521 |
| **ZOTU148** | **3.317** | **1.159** | **0.016** |
| **ZOTU70** | **3.543** | **1.345** | **0.028** |
| **ZOTU33** | **3.656** | **1.019** | **0.002** |
| **ZOTU61** | **3.981** | **1.659** | **0.046** |
| **ZOTU12** | **4.108** | **0.850** | **0.000** |
| **ZOTU105** | **4.547** | **1.530** | **0.012** |
| **ZOTU50** | **4.642** | **1.071** | **0.000** |
| **ZOTU166** | **5.235** | **1.108** | **0.000** |
| **ZOTU6** | **5.255** | **1.087** | **0.000** |
| **ZOTU155** | **5.408** | **0.735** | **0.000** |
| **ZOTU68** | **5.463** | **1.932** | **0.017** |
| **ZOTU40** | **6.825** | **1.635** | **0.000** |
| **ZOTU36** | **6.891** | **1.067** | **2E-09** |
| **ZOTU19** | **7.261** | **1.004** | **2E-11** |
| **ZOTU52** | **8.021** | **1.798** | **6E-05** |

**Table S4.** Summary of the correlations between differentially abundant ZOTUs in fertilized mixed soils and plant response variables. The response variables were total plant biomass (g dry matter) and total N content of the leaves (mg g^-1^). Perason’s correlations were measured on the relative abundances of ZOTUs (*Mesorhizobium*) present in fertilized plants, at a significance of p ≤ 0.05.

| *Taxon* | *Plant biomass (g)* | | *Total N content (mg g^-1^)* | |
| --- | --- | --- | --- | --- |
|  | *r* | *P-value* | *r* | *P-value* |
| ZOTU19 | -0.117 | 0.184 | -0.121 | 0.172 |
| ZOTU21 | 0.063 | 0.475 | 0.001 | 0.992 |
| ZOTU28 | -0.125 | 0.155 | -0.141 | 0.109 |
| ZOTU40 | -0.014 | 0.875 | 0.019 | 0.833 |
| ZOTU68 | 0.051 | 0.563 | 0.081 | 0.357 |
| ZOTU12 | 0.094 | 0.288 | 0.086 | 0.331 |
| ZOTU43 | -0.152 | 0.083 | -0.142 | 0.106 |
| ZOTU93 | 0.099 | 0.265 | 0.104 | 0.240 |
| ZOTU33 | 0.080 | 0.367 | 0.092 | 0.296 |
| ZOTU34 | -0.081 | 0.361 | -0.052 | 0.554 |
| ZOTU61 | 0.119 | 0.179 | 0.151 | 0.086 |
| ZOTU14 | 0.040 | 0.652 | 0.016 | 0.860 |
| ZOTU49 | 0.044 | 0.619 | 0.064 | 0.470 |
| ZOTU36 | -0.070 | 0.427 | 0.022 | 0.805 |
| ZOTU166 | 0.119 | 0.177 | 0.019 | 0.600 |
| ZOTU164 | 0.019 | 0.830 | 0.115 | 0.194 |
| ZOTU17 | -0.042 | 0.632 | -0.068 | 0.442 |
| ZOTU6 | -0.038 | 0.666 | 0.082 | 0.357 |
| ZOTU23 | 0.008 | 0.932 | -0.017 | 0.843 |
| ZOTU173 | 0.134 | 0.130 | 0.125 | 0.158 |
| ZOTU52 | -0.058 | 0.514 | -0.082 | 0.351 |
| ZOTU155 | -0.004 | 0.963 | -0.061 | 0.493 |
| ZOTU50 | -0.037 | 0.676 | 0.054 | 0.543 |
| ZOTU105 | -0.096 | 0.280 | -0.128 | 0.147 |
| ZOTU70 | -0.065 | 0.435 | -0.061 | 0.489 |
| ZOTU148 | 0.133 | 0.131 | 0.171 | 0.051 |

**Methods S1 Library preparation for sequencing of the *nodA* (Illumina MiSeq) amplicons**

After dilution of the *nodA* amplicons to 1 ng µl^-1^, unique Nextera XT sample indices (Illumina, San Diego, CA, USA) were added by a second PCR in a reaction volume of 25 µl with 12.5 µl KAPA HiFi Hot Star Ready Mix (KAPA Biosystems Inc., Wilmington, MA, USA), 5 µl PCR-grade water, 2.5 µl of each of the index primers at a concentration of 10 µM and 2.5 µl of the purified amplicons as template. The thermal cycling program was the following: initial denaturation at 95°C for 3 min, 8 cycles of denaturation at 95°C, primer annealing at 55°C and primer extension at 72°C for 30 s and a final further extension at 72°C for 5 min. The indexed PCR products were bead-purified using the same protocol as for the initial amplicons and quantified using the Qubit assay (Thermo Fisher Scientific Inc.) on a Spark 10M Multimode Plate Reader (Tecan). The *nodA* amplicon size and integrity was verified on the Agilent 2200 Tape Station (Agilent, USA) before dilution to 4 ng μl^-1^ for equimolar pooling of all 266 samples for library preparation. The amplicons were 2x300 bp paired-end sequenced, using the Illumina MiSeq v2 chemistry and PhiX as internal standard at a concentration of 48.99% on an Illumina MiSeq sequencer at the Genetic Diversity Center of ETH Zürich, Switzerland.

**Methods S2 Bioinformatical analyses on *gyrB* and *nodA* sequencing data**

The unique *nodA* sequence reads were obtained with the dereplication command *fastx_uniques* in USEARCH and Illumina sequencing errors were corrected and chimeras removed in UNOISE3. ZOTUs were determined using the same approach as for the *gyrB*. The count table was generated, using the *otutab* command in USEARCH. The phylotaxonomic affiliations of the ZOTUs and a further chimera check were done in SINTAX (Edgar, 2016) in comparison to all *nodA* sequences on NCBI GenBank.
